# Reconstruction of ancestral genomes in presence of gene gain and loss

**DOI:** 10.1101/040196

**Authors:** Pavel Avdeyev, Shuai Jiang, Sergey Aganezov, Fei Hu, Max A. Alekseyev

## Abstract

Since most dramatic genomic changes are caused by genome rearrangements as well as gene duplications and gain/loss events, it becomes crucial to understand their mechanisms and reconstruct ancestral genomes of the given genomes. This problem was shown to be NP-complete even in the “simplest” case of three genomes, thus calling for heuristic rather than exact algorithmic solutions. At the same time, a larger number of input genomes may actually simplify the problem in practice as it was earlier illustrated with MGRA, a state-of-the-art software tool for reconstruction of ancestral genomes of multiple genomes.

One of the key obstacles for MGRA and other similar tools is presence of breakpoint reuses when the same breakpoint region is broken by several different genome rearrangements in the course of evolution. Furthermore, such tools are often limited to genomes composed of the same genes with each gene present in a single copy in every genome. This limitation makes these tools inapplicable for many biological datasets and degrades the resolution of ancestral reconstructions in diverse datasets.

We address these deficiencies by extending the MGRA algorithm to genomes with unequal gene contents. The developed next-generation tool MGRA2 can handle gene gain/loss events and shares the ability of MGRA to reconstruct ancestral genomes uniquely in the case of limited breakpoint reuse. Furthermore, MGRA2 employs a number of novel heuristics to cope with higher breakpoint reuse and process datasets inaccessible for MGRA. In practical experiments, MGRA2 shows superior performance for simulated and real genomes as compared to other ancestral genomes reconstruction tools. The MGRA2 tool is distributed as an open-source software and can be downloaded from GitHub repository http://github.com/ablab/mgra/. It is also available in the form of a web-server at http://mgra.cblab.org, which makes it readily accessible for inexperienced users.

## 1 Introduction

Recent advances in high-throughput sequencing and the rapidly growing number of assembled genomes emphasizes the need for new algorithms to analyze the genomes and extract phylogenomic information about large genetic variations. Since most dramatic genomic changes are caused by *genome rearrangements* (such as *reversals*, *translocations*, *fissions*, *fusions*) as well as gene *duplications* and *gain/loss* events, it becomes extremely important to understand their mechanisms and reconstruct the common ancestral genomes and sequence of evolutionary events (*evolutionary history*) between genomes of interest.

The problem of reconstruction of ancestral genomes from genomes of living species (with a known phylogenetic tree) has a rich history of research. The GRAPPA algorithm (Moret et al., 2001) computes the reversal distance between unichromosomal genomes of a given phylogeny tree. The inferCARs algorithm (Ma et al., 2006) recovers contiguous ancestral regions based on the analysis of gene adjacencies. The EMRAE algorithm (Zhao and Bourque, 2007) identifies reliable genome rearrangements over conserved gene adjacencies across extant genomes. The MGR tool (Bourque and Pevzner, 2002) implements an algorithm, which minimizes the total number of rearrangements over all the branches of the phylogenetic tree. It was further improved in MGRA (Alekseyev and Pevzner, 2009), which is able to reveal rearrangements in ancestral genomes even before they are reconstructed. InferCARsPro (Ma, 2010) and PMAG (Hu et al., 2013) are probabilistic homology-based methods that compute the probability of each gene adjacency and reconstruct the ancestral genomes based on these probabilities.

The aforementioned methods are restricted to genomes with equal gene content, when each gene is present in every genome in exactly one copy. In reality, different genes may be missing in different lineages. In fact, as the number of genomes grows, the number of genes shared across all these genomes decreases substantially (Peng et al., 2010). On the other hand, genomes that share more genes are likely to be evolutionarily closer to each other. Thus, revealing evolutionary insertions and deletions (*indels*) of genes becomes crucial for understanding relationship between multiple extant genomes.

While some studies include indels in the rearrangement analysis (Yancopoulos and Friedberg, 2008; Bader, 2009; Braga et al., 2011; Compeau, 2013; Willing et al., 2013), they are mostly of theoretical nature and/or limited to the case of two genomes. We are aware of the following software tools that can reconstruct ancestral genomes for input genomes with unequal gene content. Egchel (Arndt and Tang, 2011) converts given genomes with unequal gene content into genomes with equal gene content by introducing *prosthetic* chromosomes that minimize the sum of pairwise rearrangement distances between the genomes. GapAdj (Gagnon et al., 2012) is a homology-based method that models indels as well as whole genome duplications in addition to rearrangements. PMAG^+^ (Hu et al., 2014) is an extension of PMAG that supports indel operations.

In the current work, we present a next-generation tool MGRA2 that not only organically incorporates indels into the rearrangement analysis of multiple genomes but also employs a number of novel heuristics to cope with higher breakpoint reuse and process datasets inaccessible for MGRA. We conduct a series of experiments to evaluate the performance of MGRA2 and compare it to GapAdj and PMAG^+^ tools on simulated and real mammalian genomes. The experimental results show that MGRA2 outperforms other ancestral genomes reconstruction tools for both simulated and real genomes.

We remark that while MGRA2 (similarly to MGRA) can be used for imposing phylogeny of input genomes, in this work we address the ancestral genomes reconstruction problem for input genomes with a known phylogenetic tree. Some methods for imposing phylogeny, such as TIBA (Lin et al., 2012) and MLWD (Lin et al., 2013), can account for rearrangements and indels but do not reconstruct ancestral genomes and thus are not used in our practical experiments.

The MGRA2 tool is distributed as an open-source software and can be downloaded from GitHub repository http://github.com/ablab/mgra/. It is also available in the form of a web-server at http://mgra.cblab.org, which makes it readily accessible for inexperienced users.

## 2 Overview of MGRA

For given genomes on the same set of genes (or, more generally, synteny blocks) and their evolutionary tree, MGRA (Alekseyev and Pevzner, 2009) reconstructs ancestral genomes at the internal nodes of the tree under the assumption that evolutionary events consist only of genome rearrangements. In this section we give an overview of the main algorithmic ideas employed in the MGRA framework.

### 2.1 Breakpoint Graphs and 2-Break Rearrangements

Let *P*_1_, *P*_2_, …, *P*_*k*_ be genomes composed of linear and/or circular chromosomes on the same set of synteny blocks. We associate with each genome *P*_*i*_ a unique color, referred to as color *P*_*i*_; an edge of color *P*_*i*_ is called a *P*_*i*_-*edge*. Yet another color called *obverse* is reserved for coloring of the blocks.

A circular chromosome in genome *P*_*i*_ consisting of *n* synteny blocks is represented as a sequence of *n* directed obverse edges of the form (*b*^*t*^, *b*^*h*^) or (*b*^*h*^, *b*^*t*^) encoding blocks and their strands, where *b*^*t*^ and *b*^*h*^ are extremities (“tail” and “head”) of block *b*, and *n* undirected *P*_*i*_-edges that connect extremities of adjacent blocks. Thus, each circular chromosome in genome *P*_*i*_ represents an alternating cycle of obverse and *P*_*i*_-edges. A linear chromosome in genome *P*_*i*_ consisting of *n* synteny blocks is similarly represented by *n* directed obverse edges, *n* – 1 undirected *P*_*i*_-edges that connect extremities of adjacent blocks, and 2 more undirected edges of color *P*_*i*_ that connect extremities of the first and last block to the special vertex ∞.^1^

We will not explicitly deal with the obverse edges (which can be recovered from vertex labels) and focus on the edges of colors *P*_1_, …, *P*_*k*_. A *genome graph G*(*P*_*i*_) consists of all edges of the *P*_*i*_ color representing the chromosomes of genome *P*_*i*_ (Fig. 1a).

**Figure 1:**
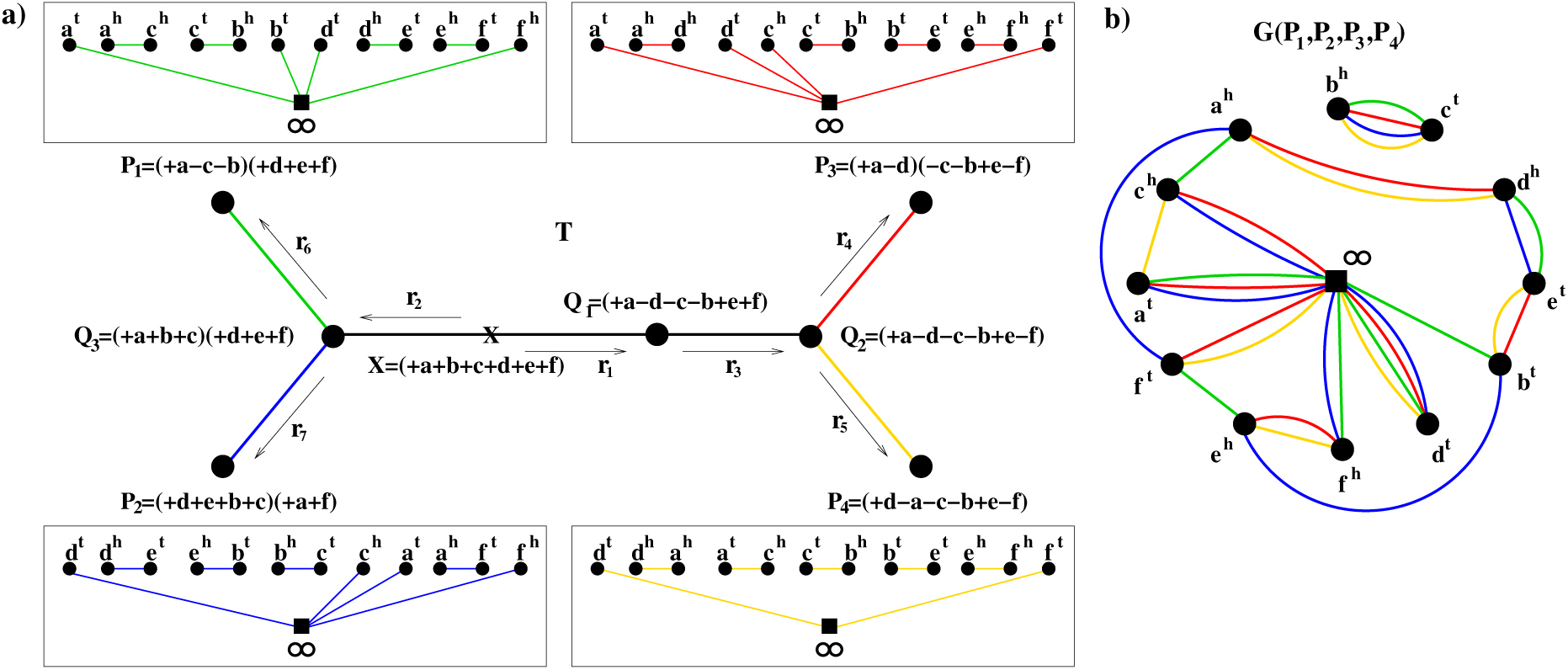
An example of four genomes with a phylogenetic tree and their multiple breakpoint graph. a) A phylogenetic tree with four genomes *P*_1_ (green), *P*_2_ (blue), *P*_3_ (red), *P*_4_ (yellow) at the leaves and specified intermediate genomes. Genomes *P*_1_, *P*_2_, *P*_3_, *P*_4_ are represented as genome graphs (the obverse edges are not shown). b) The breakpoint graph *G*(*P*_1_, *P*_2_, *P*_3_, *P*_4_) is the superposition of the genome graphs.

A *2-break* (Alekseyev and Pevzner, 2008; Alekseyev, 2008) (first introduced as DCJ by (Yancopoulos et al., 2005)) in genome *P*_*i*_ replaces a pair of *P*_*i*_-edges with another pair of *P*_*i*_-edges forming a matching on the same four vertices [e.g., in Fig. 1b a possible 2-break can replace a pair of blue edges (*a*^*h*^, *ƒ*^*t*^) and (*b*^*t*^, *e*^*h*^) with a pair of blue edges (*a*^*h*^, *b*^*t*^) and (*ƒ*^*t*^, *e*^*h*^), where each pair forms a matching on the same vertices *a*^*h*^, *ƒ*^*t*^, *b*^*t*^, *e*^*h*^]. Reversals, translocations, fissions, fusions in a genome represent all different types of 2-breaks in its genome graph.

The *breakpoint graph G*(*P*_1_, …, *P*_*k*_) is defined as the superposition of individual genome graphs *G*(*P*_1_),*G*(*P*_2_), …, *G*(*P*_*k*_) and can be constructed by “gluing” the identically labeled vertices in the genome graphs. The breakpoint graph consists of undirected edges of *k* colors *P*_1_, …, *P*_*k*_ representing block adjacencies in the corresponding genomes (Fig. 1b).

### 2.2 MGRA Framework

Let 𝒞 be the set of all colors {*P*_1_, …, *P*_*k*_}, a non-empty subset of 𝒞 is called *multicolor*. All edges connecting vertices *x* and *y* in the breakpoint graph *G*(*P*_1_, …, *P*_*k*_) form the *multiedge* (*x, y*) of the multicolor composed of the colors of these edges. The number of multiedges connected to a vertex *x* is called *multidegree* of *x*. A vertex is called *breakpoint* if its multi-degree is greater than 1. A breakpoint graph without breakpoints is an *identity breakpoint graph G*(*R*, *R*, …, *R*) of some genome *R*. Alternatively, the identity breakpoint graph can be characterized as a breakpoint graph consisting of *complete multiedges* of multicolor 𝒞 [e.g., in Fig. 1b multiedge (*b*^*h*^, *c*^*t*^) is complete], which correspond to the synteny block adjacencies in *R*.

Let *X* be a common ancestral genome of genomes *P*_1_, …, *P*_*k*_, then each genome *P*_*i*_ was obtained from *X* with a series of rearrangements. Therefore, there exists a transformation of *k*-tuples of genomes and their breakpoint graphs:

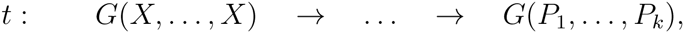

where each step represents a single 2-break occurring in either in one of genomes *P*_1_, …, *P*_*k*_ (in which case only one genome in the tuple changes) or in some of the ancestral genomes (in which case multiple genomes sharing this ancestor change simultaneously). In other words, each 2-break in this transformation occurs either in a single color or in a multicolor corresponding to an ancestral genome. Such multicolors are called *T-consistent* and formed by the colors of the leaves when they are partitioned by removal of a branch from *T*. Therefore, the (undirected/directed) branches of *T* are in one-to-one correspondence with (unordered/ordered) pairs of complementary *T*-consistent multicolors (Supplementary Fig. S1). If multicolor *Q* is *T*-consistent, then so is 𝒞\*Q*, i.e., *T*-consistent multicolors form pairs of complementary multicolors.

We remark that transformation *t* can be viewed as a collection of paths in *T*, starting at the genome *X* and ending at each of the leaf genomes. Every internal node of *T* is visited by some of these paths and they define a genome at this node. Therefore, to reconstruct the genomes at internal nodes of *T*, it is enough to find *t*. MGRA attempts to reconstruct a reverse transformation for *t*:

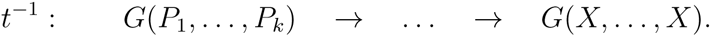

We remark that the property of *X* being a common ancestral genome of *P*_1_, …, *P*_*k*_ is inessential, *X* can be any genome from the course of their evolution (i.e., *X* resides in *T*), including the genomes *P*_1_, …, *P*_*k*_ themselves. We therefore do not fix genome *X* but determine it *post factum*. In other words, MGRA attempts to find a transformation of *G*(*P*_1_, …, *P*_*k*_) into *any* identity breakpoint graph with the minimum number of 2-breaks. Alternatively, this transformation can be viewed as elimination of breakpoints in *G*(*P*_1_, …, *P*_*k*_).

We remark that in transformation *t*, all 2-breaks are “directed” from *X* towards the leaf genomes, implying that in any pair of complementary *T*-consistent multicolors, only one multicolor may be used by 2-breaks of *t* (the only exception is the multicolors corresponding to the branch where *X* resides). To enforce this restriction, MGRA fixes a branch *𝒳* of *T* assuming that *X* resides on *𝒳*, breaks *𝒳* into two subbranches, and directs all branches of *T* toward *X* (Supplementary Fig. S1). Multicolors that correspond to the starting nodes of each directed branch are called 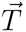-*consistent*^2^ and only they are allowed for use by 2-breaks in *t*^−1^. For example, in the tree *T* in Supplementary Fig. S1, the multicolor *MR* is 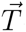-*consistent*, while *RDQHC* is *T*-consistent but not 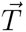-consistent.

Another important property of *t*^−1^ (and *t*) is that it must also be a *strict* transformation, i.e., for each pair of 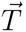-consistent multicolors *Q*_1_ ⊊ *Q*_2_, all 2-breaks in *Q*_1_ must precede all 2-breaks in *Q*_2_ in *t*^−1^ (respectively, in *t* all 2-breaks in *Q*_2_ must precede all 2-breaks in *Q*_1_). However, having a transformation *t*′ of *G*(*P*_1_, …, *P*_*k*_) into *G*(*X*, …, *X*) with 2-breaks in 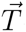-consistent multicolors in any order, it is possible to reorder 2-breaks in this transformation to obtain a strict transformation.^3^ Therefore, MGRA does not initially enforce strictness and constructs a transformation *t*′, then reorders *t*′ into *t*^−1^, and eventually recovers *t* and the ancestral genomes.

To construct *t*′, the MGRA involves a number of stages based on exact and heuristic genome sorting algorithms. More reliable stages (based on exact algorithms) come first, while less reliable heuristics come last. The number of MGRA stages to perform can be limited by a user, in which case MGRA may not be able to complete the transformation *t*′ required for reconstruction of the ancestral genomes. However, in this case by placing the target genome *X* at a particular internal node of *T*, MGRA can reliably reconstruct Conserved Ancestral Regions (CARs) of the ancestral genome at this node. Below we describe major design principles and stages of rearrangement analysis within the MGRA framework.

### 2.3 MGRA Rearrangement Analysis

As explained above, the first principle of MGRA is:

(P1) MGRA performs 2-breaks only in 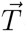-consistent multicolors.

MGRA relies on some observations from the well-studied rearrangement analysis of two genomes and extends them to the case of multiple genomes. In particular, it is easy to see that the breakpoint graph of two genomes that are one 2-break apart contains a single non-trivial component, namely a cycle on four vertices and four edges. It is possible to identify these vertices and revert the 2-break, even if we observe only a path on three of these edges. MGRA uses the cycles/paths in the breakpoint graphs as a guidance for finding reliable rearrangements. A vertex (breakpoint) is *simple* if its multidegree is 2. A path/cycle is called *good* if all the internal vertices are simple and the multiedges connecting these vertices alternate between 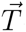-consistent multicolor *Q* and its complement 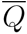 [e.g., in Fig. 1b multiedges (*a*^*h*^, *d*^*h*^), (*d*^*h*^, *e*^*t*^), and (*e*^*t*^, *b*^*t*^) form a good path with multicolors alternating between red+yellow and green+blue]. A good cycle on 2*m* multiedges alternating between 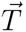-consistent multicolor *Q* and its complement 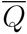 is transformed into *m* complete multiedges with *m*-1 2-breaks in multicolor *Q*. Similarly, a good path with *m* multiedges of 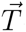-consistent multicolor *Q* is first transformed with a 2-break on its first and last multiedges of multicolor *Q* into a good cycle, which is then transformed into *m* - 1 complete multiedges with *m* - 2 2-breaks.

It is easy to see that 2-breaks processing of good paths/cycles do not decrease the number of connected components. This is a particular case of more general principle:

(P2) MGRA never decreases the number of connected components in the breakpoint graph.

This principle is inspired by the following property of shortest rearrangement scenarios of two genomes. While the breakpoint graph of two genomes is formed by paths and cycles, representing its connected components, any identity breakpoint graph consists of complete multiedges, each of which forms its own connected component. A transformation of the breakpoint graph into an identity breakpoint graph with 2-breaks thus can be viewed as a process of increasing (but never decreasing) the number of connected components.

*Breakpoint reuses* (Alekseyev, 2008; Alekseyev and Pevzner, 2010) occur when different 2-breaks operate on the same vertices. As a result, the breakpoint graph may contain *complex breakpoints* (i.e., vertices of multidegree *>* 2) and multiedges of non-*T*-consistent multicolors. While MGRA is not allowed to move such multiedges, it can enrich their multicolors by making other multiedges parallel, possibly transforming them into *T*-consistent or even complete multiedges. If this has not happened, at later stages MGRA can also break non-*T*-consistent multiedges into *T*-consistent submultiedges and perform 2-breaks on such submultiedges. However, MGRA believes that *T*-consistent multiedges correspond to adjacencies in the corresponding ancestral genomes and thus:

(P3) MGRA never breaks *T*-consistent multicolors into submulticolors.

This principle may be viewed as a generalization of so-called *perfect rearrangement scenarios* (Bérard et al., 2009) to the case of multiple genomes.

We observed that the most common case of complex breakpoints appears in a form of *composite* multiedges (*x, y*), where both *x* and *y* have multidegree of 3 and there exists a one-to-one correspondence between the multiedges incident to *x* and *y* that is invariant with respect to the multicolor [e.g., in Fig. 1 a multiedge (*ƒ*^*1*^, *e*^*h*^) is composite as vertices *ƒ*^*t*^ and *e*^*h*^ have incident multiedges of the same multicolors]. A composite multiedge (*x, y*) is called *fair* if all multiedges incident to *x* and *y*, except (*x, y*), are 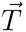-consistent, while (*x, y*) is not 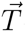-consistent. Fair edge (*x, y*) is transformed into a complete multiedge with two 2-breaks on the matching 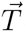-consistent edges incident to *x* and *y*. We remark that result of these 2-breaks do not depend on their order.

## 3 MGRA2 Framework

### 3.1 Indels Analysis in MGRA2

When a block *b* is present in all genomes *P*_1_, …, *P*_*k*_, the corresponding vertices *b*^*t*^ and *b*^*h*^ in *G*(*P*_1_, …, *P*_*k*_) are incident to edges of all *k* genome colors. On the other hand, when *b* is absent in genome *P*_*i*_, the vertices *b*^*t*^ and *b*^*h*^ have no incident edges of *P*_*i*_-color. A vertex in *G*(*P*_1_, …, *P*_*k*_) is called *unbalanced* if the number of its incident edges is less than *k* (i.e., some of the genome colors are absent in the incident edges). We remark that for each unbalanced vertex, its obverse counterpart vertex is also unbalanced, and they miss the same colors among their incident edges.

Insertion and deletion events result in unequal block contents across the genomes, which further imply appearance of unbalanced vertices in the breakpoint graph. To incorporate processing of indels into the MGRA2 framework, we generalize the idea of *prosthetic chromosomes* (Arndt and Tang, 2011; Compeau, 2013) used in studies of indels between two genomes. Namely, for a pair of unbalanced obverse vertices *b*^*t*^ and *b*^*h*^, whose missing incident edges’ colors form a set *Q* (in other words, *Q* is the set of genomes that lack block *b*), we balance them by adding in the breakpoint graph a *prosthetic multiedge* (*b*^*t*^, *b*^*h*^) of the multicolor *Q* (Fig. 2a). While this multiedge may be viewed as a circular chromosome on a single block *b* in each of the genomes from *Q*, we do not assume any genomic interpretation for this multiedge as itself. Instead, we view it as a “dark hole” which may supply edges for insertion or absorb edges being deleted.

**Figure 2:**
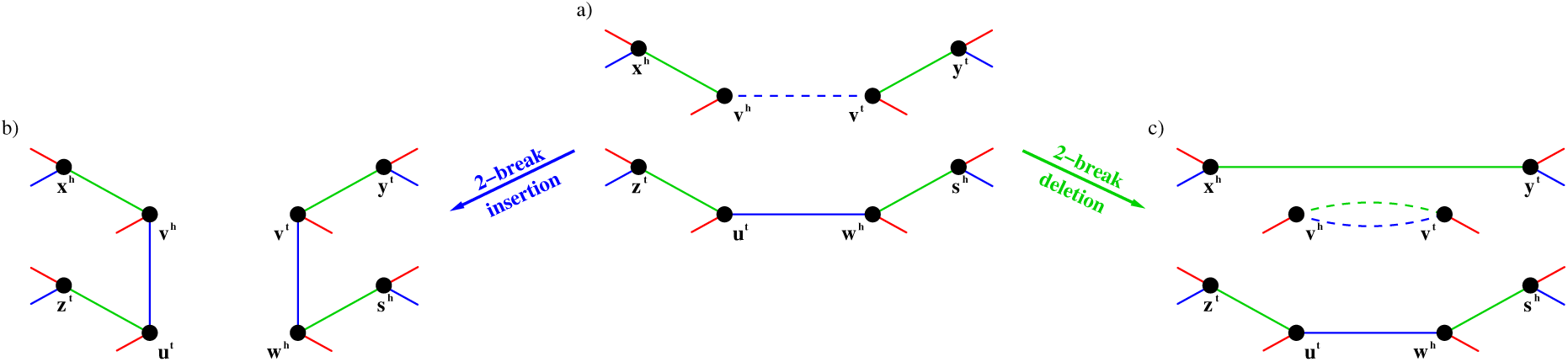
a) A subgraph of the balanced breakpoint graph of three genomes colored blue, red, and green. The blue edge (*w*^*h*^, *u*^*t*^) encodes adjacent blocks *wu* in the blue genome, while the blue prosthetic edge (*v*^*t*^, *v*^*h*^) mimics the absence of block *v*. b) The subgraph after a 2-break in the blue genome that replaces the prosthetic edge (*v*^*t*^, *v*^*h*^) and the regular edge (*w*^*h*^, *u*^*t*^) with regular edges (*w*^*h*^, *v*^*t*^) and (*v*^*h*^, *u*^*t*^), which encode consecutive blocks *wvu*, i.e., previously absent block *v* gets inserted in between of blocks *w* and *u*. c) The subgraph after a 2-break in green genome that replaces regular edges (*x*^*h*^, *v*^*h*^) and (*v*^*t*^, *y*^*t*^), which encode consecutive blocks *x* - *vy*, with edges (*v*^*h*^, *v*^*t*^) and (*x*^*h*^, *y*^*t*^). The parallel green and blue edges (*v*^*h*^, *v*^*t*^) form a prosthetic multiedge, while the green edge (*x*^*h*^, *y*^*t*^) encodes adjacent blocks *xy*, i.e., block *v* gets deleted from the green genome.

After balancing, the graph undergoes to rearrangement analysis within the MGRA2 framework, which makes no distinction between regular and prosthetic multiedges. However, in the resulting transformation, we interpret 2-breaks that involve prosthetic multiedges as insertions or deletions. Namely, a 2-break that moves a prosthetic (sub)multiedge of multicolor *Q* to a different position is interpreted as an insertion in *Q* (Fig. 2a,b), while a 2-break that moves a regular edge of multicolor *Q* into a prosthetic position is interpreted as a deletion in *Q* (Fig. 2a,c). We further remove from the resulting identity breakpoint graph all complete multiedges in the prosthetic positions. This way, we obtain a transformation of the original unbalanced breakpoint graph (with no prosthetic multiedges) that involves both rearrangements and indels and may result in an identity breakpoint graph where some vertices are absent (such vertices correspond to deleted blocks in the underlying genome). We remark that the total number of rearrangements and indels equals the number of rearrangements in the original transformation involving prosthetic edges.

### 3.2 Rearrangement Analysis in MGRA2

While MGRA2 inherits the overall framework and principles (P1)-(P3) from MGRA, its rearrangement analysis is improved and can handle situations beyond the reach of MGRA. In particular, in most cases MGRA2 restricts its attention to a small subgraph of the breakpoint graph to find reliable rearrangements (*locality* property) and relies on a new notion of *multiedge mobility* that generalizes the notion of 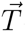-consistency and extends its applicability from good paths/cycles and fair edges to more complex features in the breakpoint graphs.

#### Locality

We remark that for any two genomes *P*_1_ and *P*_2_ on the same set of blocks, there exists a shortest transformation of *P*_1_ into *P*_2_ that consists of 2-breaks that always operate on two *P*_1_-edges connected by a *P*_2_-edge [e.g., *P*_1_-edges (*x*_1_, *y*_1_) and (*x*_2_, *y*_2_) such that (*x*_1_, *x*_2_) is a *P*_2_-edge]. In other words, there exists a transformation of *G*(*P*_1_, *P*_2_) into an identity breakpoint graph (with 2-breaks in both *P*_1_ and *P*_2_) where each 2-break operates on two edges that are connected by another edge. This easily generalizes to processing of good paths/cycles in the breakpoint graph of multiple genomes and further inspires the following principle:

(P4) MGRA2 performs 2-breaks on two multiedges only if they are connected by another multiedge.

We remark that this principle generalizes principle (P2) of MGRA. According to principle (P4), to find reliable rearrangements involving a particular multiedge *e*, it is enough to explore just a small neighborhood of *e* (namely, within paths of length 3 starting at *e*).

#### Multiedge mobility

We observe that in a transformation of the breakpoint graph into an identity breakpoint graph with 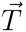-consistent 2-breaks, the multicolors of multiedges incident to any fixed vertex may change as follows. Each 2-break in the transformation either does not affect these multicolors, or unite some two of them into a single multicolor. In the latter case, the 2-break operates in one of multicolors being united and therefore this multicolor is 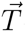-consistent. The following definition characterizes what multicolors may potentially emerge this way. A set of disjoint multicolors {*Q*_1_, *Q*_2_, …, *Q*_*m*_} is called *collapsible* if it can be collapsed into a single multicolor *Q*_1_ ∪ *Q*_2_ ∪ … ∪ *Q*_*m*_ with *m* - 1 steps, where each step unites two multicolors, at least one of which is 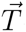-consistent, into a single multicolor.

Principles (P1) and (P4) imply that a multiedge (*x, y*) of multicolor *Q* may be moved only if *Q* is 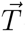-consistent and there is a multiedge (*z, t*) of multicolor *Q* such that (*x, z*) or (*y, z*) is also a multiedge. While such multiedge (*z, t*) may not be immediately available, we have some power to check whether it may become available later, inspiring the following definition.

A multiedge (*x, y*) of multicolor *Q* is called *mobile* if *Q* is 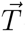-consistent and there exists a vertex *z* adjacent to *x* or *y* (*z* ≠ *x*, *z* ≠ *y*) with incident multiedges of multicolors *S*_1_, *S*_2_, …, *S*_*m*_ that are collapsible and *S*_1_ ∪ *S*_2_ ∪ … ∪ *S*_*m*_ = *Q*. The number of such vertices *z* is called *mobility* of the multiedge (*x, y*) [e.g., in Fig. 1b a multiedge (*d*^*h*^, *e*^*t*^) of green+blue multicolor has mobility 2 since there are adjacent vertices *a*^*h*^ and *b*^*t*^ where formation of incident green+blue multiedges is possible]. In other words, mobility of (*x, y*) equals the number of multiedges (currently existing or possibly appearing later) that together with (*x, y*) form a pair suitable for a 2-break. So mobile multiedges have positive mobility, while nonmobile multiedges have zero mobility.

We are particularly interested in nonmobile multiedges, since they cannot be moved by a 2-break until their multicolors are enriched by other multiedges (which may make them mobile). In fact, in many cases nonmobile multiedges remain throughout the whole transformation and thus identify positions of complete multiedges in the resulting identity breakpoint graph. Clearly, each non-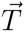-consistent multiedge is nonmobile but some 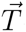-consistent multiedges may also be nonmobile. This extends applicability of many algorithms within MGRA2 that rely on the fact that a multiedge cannot move.

In MGRA2, we generalize the notion of simple paths and fair edges as follows. A nonmobile multiedge (*x, y*) of multicolor *Q* is called *fixed* if all multiedges incident to *x* and *y* are 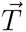-consistent and there exists a one-to-one correspondence between the multiedges incident to *x* and *y* that is invariant with respect to the multicolor. Fixed edges can be transformed into complete multiedges with 2-breaks on the matching 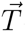-consistent multiedges incident to *x* and *y*. Clearly, fair edges represent a particular case of fixed edges (with multidegree of *x* and *y* equal 3). Similarly, a good path/cycle represents a sequence of fixed multiedges (with multidegree of endpoints equal 2). MGRA2 therefore does not look for fair edges or simple path/cycles but processes them within the fixed multiedges framework, which is well consistent with principle (P4).

Processing of fixed multiedges is further generalized to the case of nonmobile multi-edges (*x, y*) with only subsets of multiedges incident to *x* and *y* being 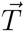-consistent and in a one-to-one correspondence. In this case, MGRA2 is guided by either the mobility of matching multiedges in these subsets or a possibility to create a *T*-consistent multiedge at (*x, y*). Namely, let these matching subsets consist of multiedges of multicolors *Q*_1_, *Q*_2_, …, *Q*_*t*_. MGRA2 performs a 2-break on the multiedges of multicolor *Q*_*i*_ only if mobility of both these multiedges equals 1 (i.e., they represents the only suitable option for each other). If no such multiedges exist, MGRA2 tries to find a subset *S* of {*Q*_1_, *Q*_2_, …, *Q*_*t*_} such the union of multicolors in *S* and multicolor of (*x, y*) is *T*-consistent, and perform 2-breaks in multicolors from *S* to turn (*x, y*) into *T*-consistent multiedge.

#### Forks and Cloning

We observe yet another common object in breakpoint graphs, called a *ƒork*, which represents an obstacle for the MGRA algorithm. A fork is formed by a *center* nonmobile multiedge (*x, y*), where *x* has multidegree 2 and *y* has multidegree 3, along with its three adjacent multiedges: *teeth* (*y, u*), (*y, v*) and a *handle* (*x, z*) of 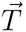-consistent multicolors *Q*_1_, *Q*_2_, and *Q* = *Q*_1_ ∪ *Q*_2_, respectively, such that mobility of the handle is 1 (Fig. 3a).

**Figure 3:**
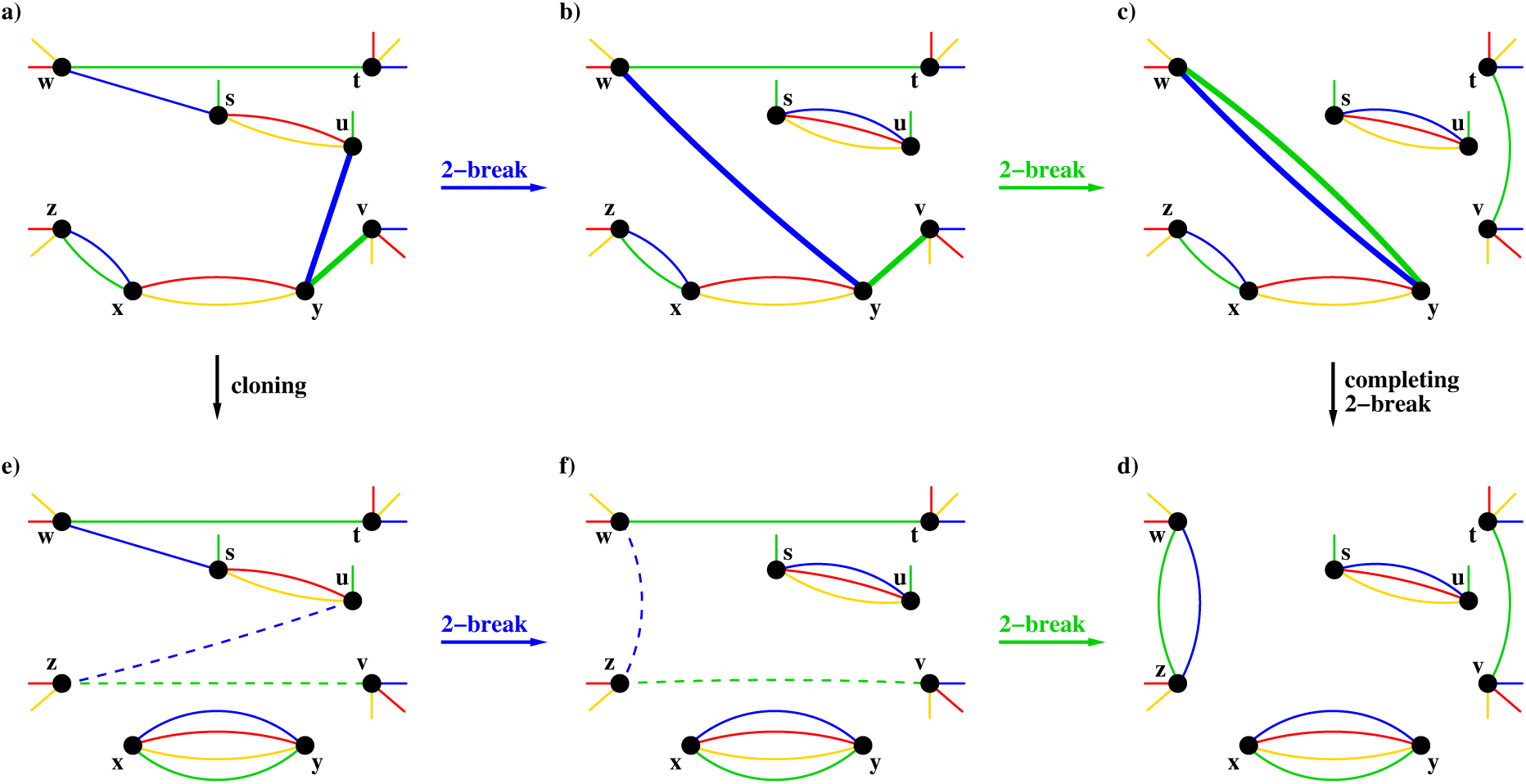
Two equivalent transformations of a breakpoint graph on red, green, blue, and yellow genomes under the assumption that the blue+green multicolor and all single colors are 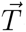-consistent, while the red+yellow multicolor is not. a) The multiedges (*z, x*), (*x, y*), (*y, u*), and (*y, v*) form a fork, whose teeth are marked by bold edges. b-c) Graphs appearing in a scenario with two 2-breaks that eventually make the fork’s teeth parallel, turning the fork into a simple path on vertices *z, x, y, w*. d) A graph after a completing 2-break that eliminates the path by creating the complete multiedge (*x, y*). e) A graph after cloning of the fork’s teeth into cloned edges (*z, u*) and (*z, v*), also resulting in the complete multiedge (*x, y*). f-d) Graphs appearing in a scenario with two 2-breaks involving cloned edges that are equivalent to the 2-breaks on regular edges in the scenario b-c. The cloned edges eventually become parallel and form a regular multiedge (*z, w*).

Since the center multiedge is nonmobile, in any transformation it stays at its place, while adjacent multiedges eventually become parallel to it and thus altogether form a complete multiedge. In other words, a fork is always a subject to the following scenario: after a number of 2-breaks the teeth converge (i.e., become parallel) and form a multiedge (*y, w*) of multicolor *Q* (Fig. 3c), which is followed by a *completing 2-break* on this multiedge and the handle (*x, z*) to form a complete multiedge at (*x, y*) (Fig. 3d). Unfortunately there may exist multiple ways for the fork’s teeth to converge and it may be not immediately possible to identify the reliable one.

In MGRA2, we propose a novel approach, called *cloning*, that creates the complete multiedge (*x, y*) in advance and lets the teeth (*y, u*) and (*y, v*) to converge afterwards. Namely, cloning replaces the teeth with the *cloned* multiedges (*z, u*) and (*z, v*) of multicolors *Q*_1_ and *Q*_2_, respectively (Fig. 3e). Any 2-break involving a cloned multiedge corresponds to a 2-break on its original multiedge, and vice versa (e.g., a 2-breaks in Fig. 3a,b is equivalent to a 2-break in Fig. 3d,e). Therefore, a cloning followed by a series of 2-breaks that make the cloned multiedges converge and result in a graph *H* corresponds to a series of 2-breaks in the original breakpoint graph (before cloning) resulting in a graph, which can be turned into *H* with a single *completing* 2-break (e.g., a scenario in Fig. 3a,e,f,d with cloning and a scenario in Fig. 3a,b,c,d without cloning are equivalent).

MGRA2 also processes generalized forks with more than two teeth and supports cloning operations of arbitrary complexity (e.g., telescopic cloning).

#### Processing Non-*T*-consistent multiedges

As we already mentioned, breakpoint reuse results in appearance of complex breakpoints (vertices of multidegree > 2) as well as multiedges of non-*T*-consistent multicolors. While breakpoint graph processing based on multiedge mobility and forks effectively eliminates complex breakpoints, non-*T*-consistent multicolors (that remain after earlier stages of MGRA2) require a special treatment.

At later stages, MGRA2 view each multiedge of non-*T*-consistent multicolor *Q* not as a single multiedge but rather as a collection of parallel submultiedges of disjoint *T*-consistent multicolors *Q*_1_, *Q*_2_, …, *Q*_*m*_ such that *Q* = *Q*_1_ ∪ … ∪ *Q*_*m*_ and *m* is minimal. Theorem 2 in Appendix A guarantees that such representation is unique.

When each non-*T*-consistent multiedge is represented as parallel *T*-consistent submultiedges (some of which are 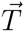-consistent per Theorem 5), the breakpoint graph can be viewed as a graph with no non-*T*-consistent multiedges. MGRA2 executes again the same algorithms searching for reliable 2-breaks (this time 2-breaks may operate on *T*-consistent submultiedges), which typically reduce the graph in significant way (in many cases simply convert it to an identity breakpoint).

As a last resort, MGRA2 has also implementation of a bruteforce approach using Blossom V software (Kolmogorov, 2009), which eliminates remaining small non-trivial connected components (to be described elsewhere).

## 4 Experiments and Evaluation

To evaluate performance of MGRA2, we run experiments on both simulated and real datasets and compare the results with PMAG^+^ (Hu et al., 2014) and GapAdj (Gagnon et al., 2012).^4^

### Simulated Genomes

We performed two sets of experiments with *N* = 6 and *N* = 12 simulated genomes. For the sake of simplicity, we limited our attention to the case of unichromosomal genomes with the evolutionary model consisting of only reversals and indels.

To simulate the genomes, we first generated a uniformly distributed rooted binary tree with *N* leaves (species). An initial unichromosomal genome with *G* = 1000 and *G* = 1500 distinct “genes” was assigned at the root. Random reversals, gene deletions and new unique gene insertions were applied to the initial genome, “moving” it along the tree branches. The total number of events along each branch was randomly chosen from the range 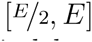 for *E* = 100 and *E* = 200. Among these events, *R* ∈ {0%, 20%, 40%, 60%} are gene indels with exactly half being gene insertions and the other half being gene deletions, while the rest are reversals. The equal number of gene insertions and deletions along each branch guarantees that the size of all genomes at the tree nodes is *G*.

For each combination of parameters *N, G, E, R*, we simulated 10 datasets and obtained simulated genomes at the leaves and internal nodes of the tree. We run MGRA2, PMAG^+^, and GapAdj on the leaf genomes and reconstructed genomes at the internal nodes. Fig. 4 reports the average accuracy measured in terms of correct (true positives), missing (false negatives), and incorrect (false positives) among gene adjacencies in the reconstructed genomes as compared to the known simulated genomes. Similarly, Fig. 5 reports the average DCJ-indel distance (computed with the (Braga et al., 2011) algorithm) in the pairs of corresponding reconstructed and simulated ancestral genomes. While GapAdj typically demonstrates the worst performance in these experiments, it is particularly sensitive to the percentage of indels with significant performance degradation for *R* > 0. In turn, PMAG^+^ is more sensitive to the number of rearrangements than the percentage of indels and performs much worse for *E* = 200 than for *E* = 100. In contrast, MGRA2 is barely sensitive to neither *R* or *E* and clearly outperforms GapAdj and PMAG^+^ in each of the metrics.

**Figure 4:**
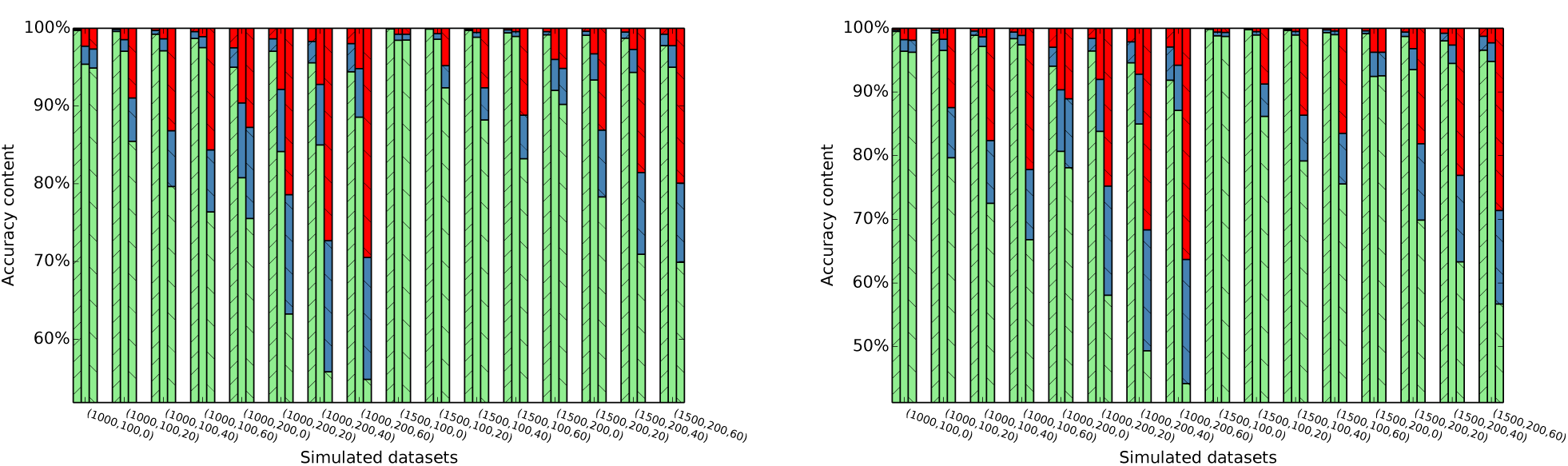
Normalized true positives (green bars), false negatives (blue bars), and false positives (red bars) among the reconstructed gene adjacencies, averaged over 10 simulated datasets with parameters (*G, E, R*) for *N* = 6 genomes (left panel) and *N* = 12 genomes (right panel), where *G* is the number of “genes”, *E* defines the interval 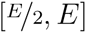 for the tree branch lengths, and *R* is the percentage of indels along each branch. For each set of parameters, the three bars correspond to MGRA2 (left bar), PMAG^+^ (middle bar), and GapAdj (right bar).

**Figure 5:**
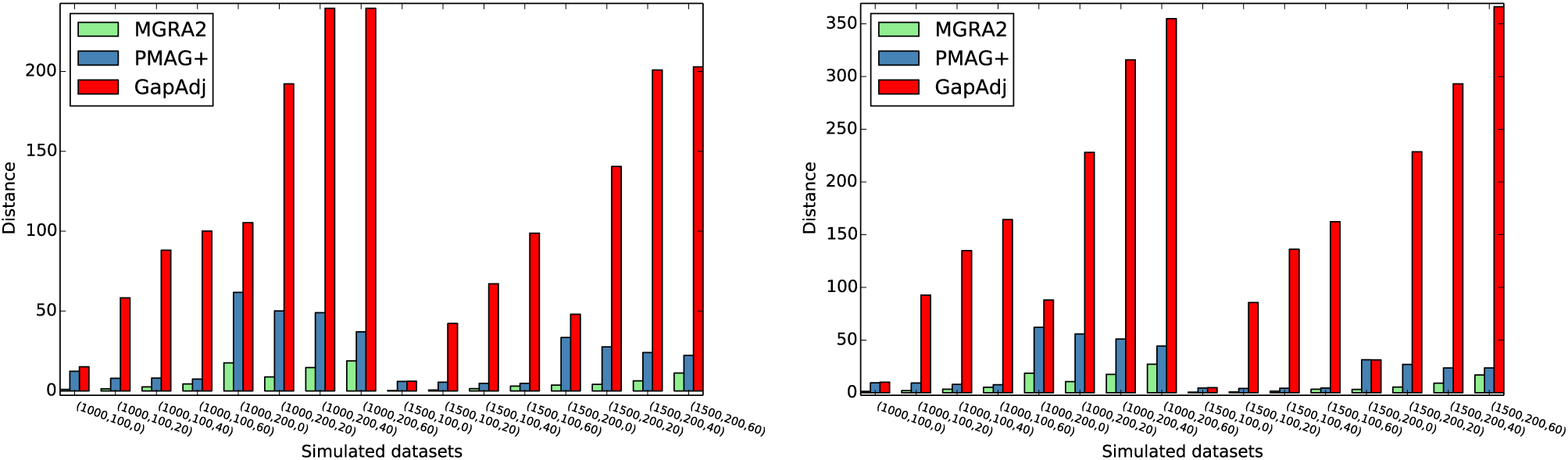
The average DCJ-indel distance between the corresponding reconstructed and simulated ancestral genomes, averaged over 10 simulated datasets with parameters (*G, E, R*) for *N* = 6 genomes (left panel) and *N* = 12 genomes (right panel), where the parameters are the same as in Fig. 4.

### Mammalian Genomes

We obtained genes of the following six mammalian genomes and their pairwise orthology relationships using Ensembl BioMart tool (Kasprzyk, 2011): *Homo sapiens* (GRCh37.p12), *Mus musculuss* (GRCm38.p1), *Rattus norvegicus* (Rnor_5.0), *Canis familiaris* (CanFam3.1), *Macaca mulatta* (MMUL.1.0), and *Pan troglodytes* (CHIMP2.1.4).

Their phylogenetic tree is shown in Fig. S1.

Since we were unable to obtain ancestral reconstructions from GapAdj on whole mammalian genomes (apparently because of their size ranging from 14, 597 genes in dog to 17, 959 genes in human), while PMAG^+^ on them produced seemingly meaningless results, we restricted this experiment only to X chromosomes (of size ranging from 591 genes in dog to 770 genes in human). The number of pairwise orthologs between X chromosomes ranged from 17, 654 (dog-chimpanzee) to 28, 732 (mouse-rat). From these pairwise orthologs we constructed 1, 009 gene families, from which we further removed those with paralogs (as they are out of scope on the current study) and obtained 879 gene families.

For each of four ancestral genomes MR (mouse-rat), MRD (mouse-rat-dog), HC (human-chimpanzee), QHC (macaque-human-chimpanzee), Fig. 6 reports unique and shared gene adjacencies across the reconstructions of X chromosome by different tools. The results show that the number of shared gene adjacencies between just MGRA2 and GapAdj is typically small (ranging from 2 to 20), which suggests that GapAdj can hardly reconstruct anything that is not reconstructed by PMAG^+^. For relatively recent ancestors MR and HC, the number of unique gene adjacencies is rather moderate for all tools, but for more ancient ancestors (MRD and QHC) this number significantly raises for GapAdj, which likely means that most of the corresponding gene adjacencies reconstructed by GapAdj are incorrect. Furthermore, for MRD and QHC genomes, MGRA2 confirms noticeable number of adjacencies (53 and 93, respectively) found by PMAG^+^ but not GapAdj, which indicates some robustness of MGRA2 and PMAG^+^ with respect to more ancient reconstructions. At the same time, for all genomes, PMAG^+^ and GapAdj share some noticeable number of gene adjacencies (ranging from 51 to 74) between themselves but not MGRA2, which possibly reflects a bias caused by the homology-based models employed PMAG^+^ and GapAdj.

**Figure 6:**
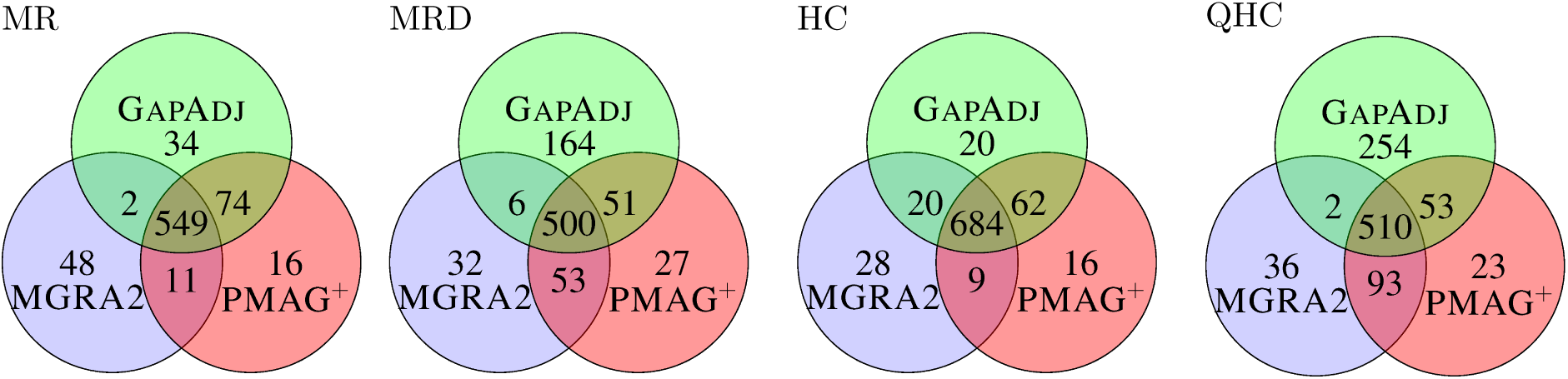
Unique and shared gene adjacencies between different reconstructions of X chromosomes of ancestral genomes MR (mouse-rat), MRD (mouse-rat-dog), HC (human-chimpanzee), and QHC (macaque-human-chimpanzee).

Table S1 gives length of the branches of the phylogenetic tree in Fig. S1 on extant and reconstructed ancestral X chromosomes. The total length for PMAG^+^ reconstructions is greater than for MGRA2 by 9%, while the total length for GapAdj reconstructions is greater by 43%. This proves that MGRA2 does a better job in minimizing the total length of the tree branches.

We also remark that in contrast to other tools, MGRA2 was able to process whole mammalian genomes and produce ancestral genomes consistent with the reconstructions of their *X* chromosomes.

## 5 Conclusion

The MGRA2 tool is a descendant of MGRA (Alekseyev and Pevzner, 2009) applicable to genomes featuring gene indels and/or high breakpoint reuse. Experiments show that MGRA2 produces significantly better results than its competitors on both simulated and real genomes. Our goals for the next release MGRA3 include fine optimization and parallelization, making it scalable to large number of genomes and large breakpoint graphs (e.g., with the locality property one can process different parts of the breakpoint graph in parallel). Another important venue for the MGRA3 development is inclusion of gene duplications (e.g., tandem duplications and whole-genome duplications) into the evolutionary model.

## Acknowledgement

The work was supported by the National Science Foundation under the grant No. IIS-1462107.

---

**Figure S1:**
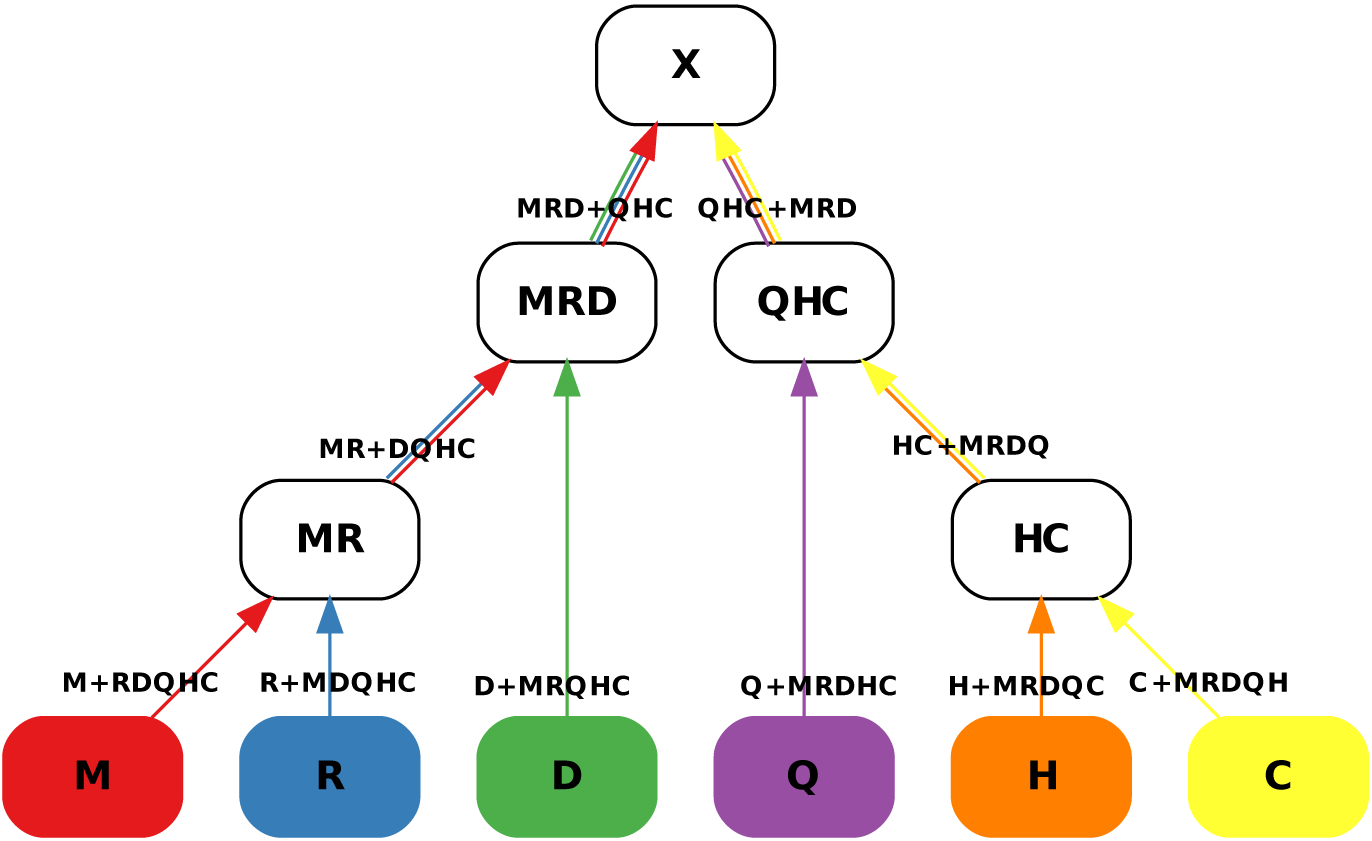
The phylogenetic tree *T* of six mammalian genomes: Mouse (*red*), Rat (*blue*), Dog (*green*), macaQue (*violet*), Human (*orange*), and Chimpanzee (*yellow*) with a root *X* on the *MRD* + *QHC* branch. The branches are directed towards *X* and labeled with the corresponding pairs of complementary *T*-consistent multicolors. The 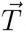-consistent multicolor from each pair also labels the starting node of the corresponding directed branch. Note that the tree orientation may not necessary correlate with the time scale and the root genome *X* may not necessarily correspond to a common ancestor of the leaf genomes. Figure is reproduced from (Alekseyev and Pevzner, 2009).

**Table S1:** Number of 2-breaks and indels (computed with the (Braga et al., 2011) algorithm) along the branches of the phylogenetic tree in Fig. S1 of mammalian X chromosomes based on the ancestral reconstructions by different tools.

| Branch | MGRA2 |  |  | GAPADJ |  |  | PMAG<sup>+</sup> |  |  |
| --- | --- | --- | --- | --- | --- | --- | --- | --- | --- |
|  | 2-breaks | indels | length | 2-breaks | indels | length | 2-breaks | indels | length |
| $MRD+$ | 0 | 47 | 47 | 38 | 156 | 194 | 24 | 65 | 89 |
| $MR+$ | 17 | 62 | 79 | 55 | 29 | 84 | 44 | 70 | 114 |
| $D+$ | 15 | 46 | 61 | 40 | 79 | 119 | 16 | 32 | 48 |
| $M+$ | 17 | 34 | 51 | 28 | 25 | 53 | 14 | 20 | 34 |
| $R+$ | 30 | 46 | 76 | 47 | 36 | 83 | 46 | 42 | 88 |
| $HC+$ | 2 | 97 | 99 | 17 | 24 | 41 | 9 | 121 | 130 |
| $Q+$ | 1 | 46 | 47 | 23 | 110 | 133 | 2 | 14 | 16 |
| $H+$ | 0 | 38 | 38 | 9 | 14 | 23 | 0 | 0 | 0 |
| $C+$ | 7 | 83 | 90 | 11 | 94 | 105 | 18 | 100 | 118 |
| Total | 89 | 499 | **588** | 268 | 567 | **835** | 173 | 464 | **637** |

## Appendix A Properties of *T* and 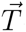-consistent multicolors

We assume that the phylogenetic tree *T* and its root *X* are fixed, which define 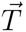-consistent and *T*-consistent multicolors composed of the leaf colors of *T*. The following lemma follows directly from the definition of *T*-consistent multicolors:

**Lemma 1**. *For any T-consistent multicolors Q*_1_ *and Q*_2_ *such that* 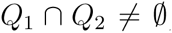, *we have Q*_1_ ⊆ *Q*_2_ *or Q*_2_ ⊆ *Q*_1_.

**Theorem 2**. *Let Q be any multicolor, then there exist a unique collection of disjoint T-consistent multicolors 𝒬* = {*Q*_1_, …, *Q*_*m*_} *such that Q* = *Q*_1_ ∪ … ∪ *Q*_*m*_ *and m is minimal*.

*Proof*. Let *R* be a *T*-consistent multicolor such that *R* ⊆ *Q* and there is no another *T*-consistent multicolor *R*′ with *R* ⊊ *R*′ ⊆ *Q*. Then *R* ∈ *𝒬* and furthermore *Q* consists of all such multicolors.

Indeed, let us consider all elements of *𝒬* that have nonempty intersection with *R*, without loss of generality, say, *Q*_1_, *Q*_2_, …, *Q*_*r*_. Since *R* ⊆ *Q*, we have *R* ⊆ *Q*_1_∪…∪*Q*_*r*_. On the other hand, by Lemma 1, each of *Q*_*i*_, 1 ≤ *i* ≤ *r*, must be a subset of *R* and thus *Q*_1_ ∪ … ∪ *Q*_*r*_ ⊆ *R*, implying that *R* = *Q*_1_ ∪ … ∪ *Q*_*r*_. Then minimality of *m* implies that *r* = 1 (otherwise replacing *Q*_1_, …, *Q*_*r*_ in *𝒬* with *R* would decrease *m*). Therefore, *R* ∈ *𝒬*.

Since for each *Q*_*i*_ ∈ *𝒬* we can find *R* (as defined above) with *Q*_*i*_ ⊆ *R*, we have *Q*_*i*_ = *R*. That is, *𝒬* entirely consists of such *R*’s.

The following lemma follows directly from the definition of *T*-consistent and 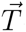-consistent multicolors:

**Lemma 3**. *Let Q be a T-consistent multicolor and let T*_*Q*_ *be the subtree oƒ induced by the leaves from Q. Then Q is* 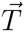-*consistent iƒ and only iƒ T*_*Q*_ *does not contain the root oƒ T*.

**Theorem 4**. *Let Q*_1_ *be a T-consistent multicolor and Q*_2_ *be a* 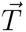-*consistent multicolor such that Q*_1_ ⊆ *Q*_2_. *Then Q*_1_ *is also* 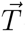-*consistent*.

*Proof*. If *Q*_1_ = *Q*_2_, the theorem statement is trivial. For the rest assume that *Q*_1_ ⊊ *Q*_2_. Let *T*′ be the subtree of *T* induced by *Q*_2_. Since *Q*_2_ is 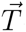-consistent, *T*′ does not contain the root of *T*.

Let *T*” be the subtree of *T* induced by *Q*_1_. Since *Q*_1_ ⊊ *Q*_2_, *T*” is a proper subtree of *T*′ and thus does not contain the root of *T*, implying that *Q*_1_ is 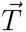-consistent.

**Theorem 5**. *Let Q be any multicolor such that Q* = *Q*_1_∪*Q*_2_∪…∪*Q*_*m*_, *where Q*_*i*_ *are disjoint T-consistent multicolors. Then all Q*_*i*_, *with possibly one exception, are* 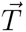-*consistent*.

*Proof*. Since *Q* is not *T*-consistent, we have *m* ≥ 2. If all *Q*_*i*_ are 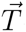-consistent, the theorem statement is proved.

If not all *Q*_*i*_ are 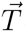-consistent, without loss of generality assume that *Q*_1_ is not 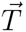-consistent. Then 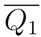 is 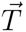-consistent. For every *i* > 1, since *Q*_1_ and *Q*_*i*_ are disjoint, we have 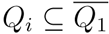, and then by Lemma 4, *Q*_*i*_ must be 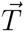-consistent.

## Footnotes

1 Everywhere below we deliberately hide details of processing the infinity vertex as in MGRA2 it is done similarly to MGRA.

2 For additional properties of 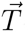-consistent and *T*-consistent multicolors, see Appendix A.

3 Such reordering is done by a stable topological sorting of 2-breaks in the transformation with respect to the partially ordered set of 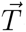-consistent multicolors.

4 For GapAdj, we used parameters *delta=25*, *threshold=0.6* as in (Hu et al., 2014); for PMAG^+^, we used the default parameters.

